# A 3D-Printed Capillary Tube Holder for High-Throughput Chemotaxis Assays

**DOI:** 10.1101/2025.08.16.670682

**Authors:** Chiara Berruto, Elisa Grillo, Shrila Esturi, Gozde S. Demirer

## Abstract

Bacterial chemotaxis is an important behavior to study to understand spatial segregation of species in mixed communities and the assembly of host microbiomes. This is particularly relevant in the rhizosphere, where chemoattraction towards root exudates is an important determinate of plant colonization. However, current methods to screen chemoeffectors are limited in their throughput, creating a barrier to generating comprehensive datasets describing chemotactic profiles for species of interest. Here, we describe a novel, 3D-printed capillary tube holder approach, which enables up to 384 simultaneous capillary tube chemotaxis assays. We optimized and benchmarked our assay using *Escherichia coli* k12 and *Bacillus subtilis* 3610 and their known chemoattractants: serine and aspartate. We then tested the threshold concentration of these chemoattractants in our assay and found we could detect chemoattraction towards concentrations spanning multiple orders of magnitudes. In this paper, we describe our novel high-throughput chemotaxis assay in detail and provide the necessary files for 3D printing the capillary tube holder.

## Introduction

Chemotaxis is the process by which motile organisms direct their movement along chemical gradients. Chemotaxis may be towards (chemoattraction) or away from (chemorepulsion) a given chemical signal facilitating the migration of cells to desirable environments. Bacteria utilize different forms of motility to carry out chemotaxis, including flagella-based swimming and pili-mediated twitching or swarming^1^. This directed motility allows bacteria to forage for nutrients, travel towards hosts or symbionts, and create spatial segregation in microbial communities^1^. This makes chemotaxis an important phenomenon to study to understand host-microbe interactions and ecological behavior.

Various tools have been developed to study bacterial chemotaxis, each having their own advantages and disadvantages. The chemotactic behavior of a bacteria in response to different chemicals can be studied using agar-based assays that involve measuring how far bacteria spread through a soft agar plate towards or away from chemoeffectors contained in hard agar plugs. The cloudy or clear zone surrounding the hard agar plug is measured to determine chemoattraction or chemorepulsion, respectively. However, this assay is low throughput allowing only a few chemoeffectors to be screened in each experiment, the measurement is dependent on the judgement of the operator, is often inconsistent^2^, and is prone to false positives^3^. Slide-based assays are an alternative strategy that involve placing cells on a microscope slide and tracking their movement towards or away from a chemoeffector using microscopy. This can be done using commercially available µ-slides and quantification can be supported by cell tracking software. While slide-based assays allow for clear visualization of cell migration and produce quantitative data, they are low-throughput and require costly equipment and software ^2^.

The capillary tube assay has, for many years, been considered the gold standard method for measuring bacterial chemotaxis. The assay involves incubating a capillary tube containing a chemoeffector in a reservoir of bacteria and measuring how many cells swim into the capillary tube via CFU counting^4^. This number is then compared to the number of cells found in a buffer control capillary tube. It is both quantitative and highly accessible for researchers, making it a popular tool to investigate bacterial chemotactic responses to different chemicals. However, the assay is low-throughput and requires precise handling of small volumes of liquids. These barriers make it challenging and time-intensive to screen many different chemoeffectors, limiting the quantity and quality of available datasets describing bacterial chemotactic profiles. A high-throughput, quantitative, reproducible, and low-cost method is still needed to enable researchers to better characterize chemotactic profiles and generate more comprehensive datasets for bacterial species of interest.

Here, we describe a novel, high-throughput, chemotaxis assay using a 3D-printed capillary tube holder that enables simultaneous screening of up to 384 chemoeffectors without the need for CFU counting. Our design is inspired by a prior approach showing that chemotaxis assays can be carried out in a plate-based fashion using lag time as a proxy for cell count^5^. We substantially improved upon this approach, replacing the Whatman UniFilter plates with a customizable 3D-printed capillary holder, effectively decreasing the cost and increasing the accessibility and throughput of the assay. We show capillary tube holder designs of a 96-well and 384-well format; however, it can easily be adapted to fit other desired plate layouts or capillary tube dimensions. To validate our method, we used previously characterized chemoattractants for *Escherichia coli* k12 and *Bacillus subtilis* 3610. The chemotactic behavior of these two species has been extensively characterized using traditional capillary tube assays, allowing us to optimize our design and benchmark our results^6,7^. We chose these two species to demonstrate that our design can be applied to both gram negative (*E. coli*) and gram positive (*B. subtilis*) species, and broadly to species with different ecological relevance.

### Design and Optimization of the 3D-Printed Capillary Tube Holder

Our 3D-printed capillary holder facilitates plate-based screening of chemoattractants and is designed to have one capillary tube per well. The dimensions of the 96-well and 384-well capillary tube holder are shown in **Figure 1A and B**, respectively. For our assays, we used Fisherbrand™ 96-Well plates and Nunc™ 384-Well plates, but the 3D-printed capillary holder is compatible with well plates of similar dimensions. The capillary holder design accommodates 1µL capillary tubes. The holder is designed to accommodate the full length of the capillary tube, holding it into the well such that the top of the capillary is flush with the top of the holder and bottom of the capillary is submerged in the plate liquid but not touching the bottom of the well. This ensures the opening to the capillary tube is accessible for the bacteria to swim freely and that the top of the capillaries can be easily sealed with a plate film. Use of capillary tubes with different dimensions will require altering the capillary holder design. See **Materials and Methods** section for the specific catalog numbers of the well plates and capillary tubes used. The capillaries must fit tightly within the capillary holder to allow for proper expulsion during the centrifugation step, which we found to be greatly influenced by the filament type used in 3D printing. Printing the capillary tube holder with thermoplastic polyurethane filament (TPU) filament improved the ability of the holder to tightly hold the capillaries without breaking compared to using polylactic acid (PLA) filament.

**Figure 1:**
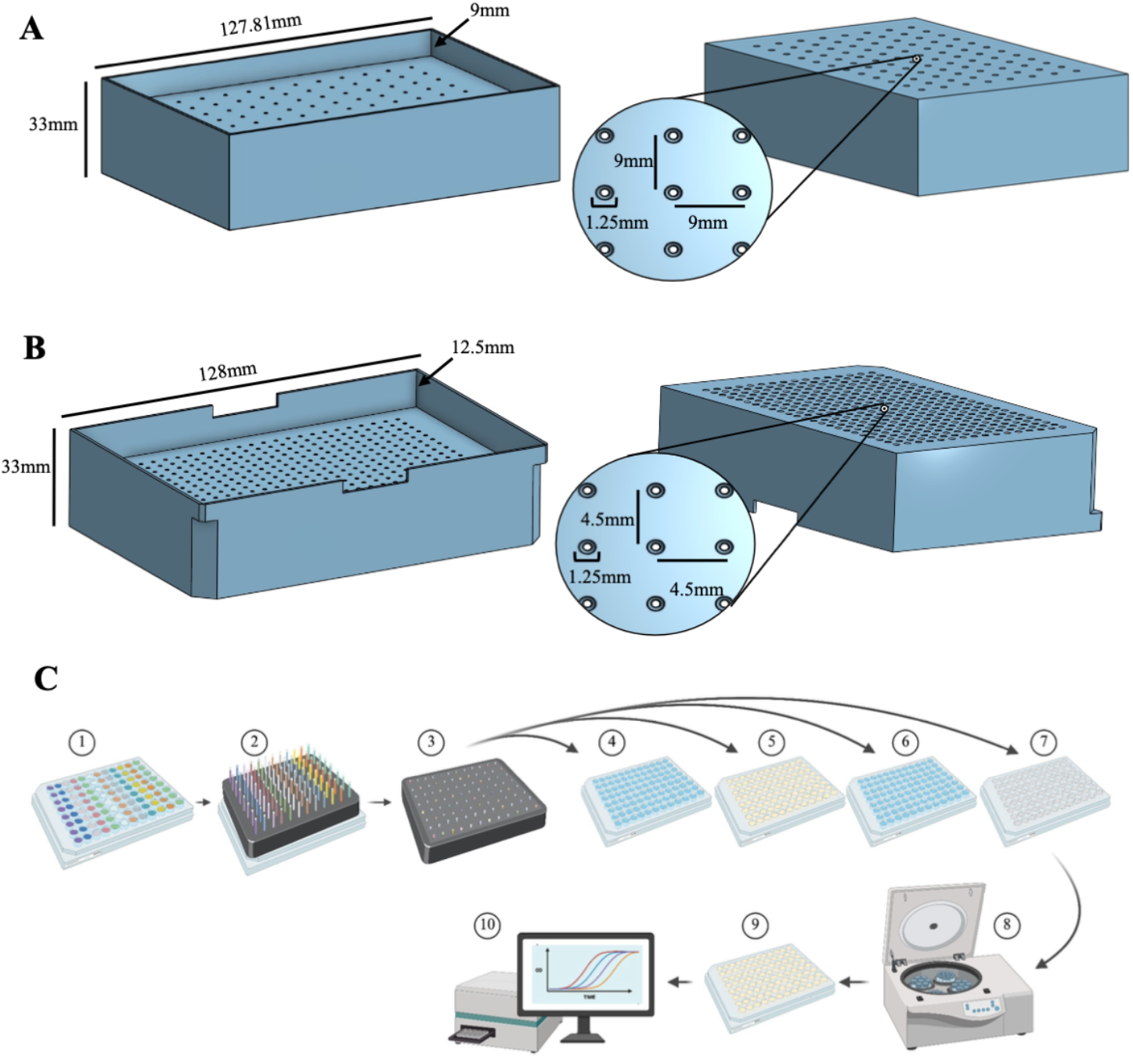
High-throughput Chemotaxis Assay Overview. Bottom (right) and top (left) view of the **A**. 96-well and **B**. 384-well capillary holder including relevant measurements. **C**. Chemotaxis assay method schematic: 1. Chemoeffector plate is filled with chemoeffectors to be tested. 2. Capillary holder is filled with capillaries and placed on top chemoeffector plate until capillaries fill with chemoeffectors. 3. Top of capillary holder is sealed with plate-sealing film. 4. Capillary holder is dipped in wash plate. 5. Capillary holder is transferred to bacteria plate and incubated for 45 min. 6. Capillary holder is dipped in wash plate. 7. Capillary holder is moved to an empty plate. 8. Capillary holder on empty plate is spun down to expel capillaries. 9. Capillary holder is removed, and growth media is added to the plate. 10. Growth curve data is collected in a plate reader.

In addition to the material used, the resolution of the 3D printer and the print conditions (see **Materials and Methods**) influence the optimal diameter of the capillary openings in the holder, and calibration may be needed for prints made under different conditions. We included a calibration print (**Supplementary File S1**), which can be printed to determine the optimal diameter for the holes in the capillary tube holder. More details on printing specifics are described in **Materials and Methods**. Design files to print the capillary tube holders and calibration piece are included as **Supplementary Files S1-S3**.

### Chemotaxis Assay Development

This method uses four well plates between which the capillary tube holder with the capillary tubes is transferred during the assay. The schematic of the method is shown in **Figure 1C**. First, the capillary tube holder is filled with capillaries and placed atop the chemoeffector plate. The capillaries fill with chemoeffectors from the plate via capillary action. The top of the holder is sealed with a plate film. The capillary tube holder with capillaries full of chemoeffectors is dipped into a wash plate containing sterile ddH_2_O water to wash the outside of the capillaries before transferring into the bacteria plate. The bacteria plate is filled with bacterial cells that are washed and resuspended in motility media, and the plate is incubated for 45 minutes to allow the bacterial cells to swim into the capillary tubes. At the end of the incubation period, the outside of the capillary tubes is washed by dipping into the wash plate and the capillary tube holder carrying the capillaries is transferred to an empty plate. The seal is removed from the top of the holder, and the capillaries are expelled into the empty plate via centrifugation. Finally, the capillary holder is removed, growth media is added to each well, and the plate is placed in a plate reader to record bacterial growth.

The data generated from the growth curve can be analyzed to record the lag time of each well, which we define as the time it takes for each well to reach an OD_600_ of 0.3. The lag time is used to calculate the initial number of cells in each well using a standard curve. From the cell counts, a chemotaxis index (CI) can be calculated using the following equation: CI = T / (T+C) where T is the number of cells in the chemoeffector capillary and C is the number of cells in the control capillary. CI values greater than 0.6 indicate an attractant, while CI values less than 0.4 indicate a repellent and CI values between 0.4 and 0.6 are considered neutral.

### Constructing Standard Curves

To quantify the number of cells in each capillary tube using growth curve data, standard curves must be constructed for each species being tested. The standard curves are used to calculate the number of cells in each well from the lag time value. To construct these standard curves, 2-fold serial dilutions of bacteria in growth media are prepared. These dilutions are CFU-counted and added to a well plate that is placed into a plate reader to measure OD_600_ overtime. From the growth curve data, the lag time can be extracted. It is critical to construct the standard curve for each species under conditions that will be used in the assay. This means preparing cells as they are prepared for the chemotaxis assay (see **Material and Methods**) and running the growth curve using the same settings and conditions that will be used when the assay is conducted.

We constructed separate standard curves for *B. subtilis* and *E. coli* in both 96-well and 384-well plates. For each standard curve, 2-fold serial dilutions were run in a plate reader; the resulting growth curves are shown in **Figure 2A**, where each line represents a different dilution of cells. As the concentration of cells gets more diluted, the lag time increases resulting in a shifted growth curve. Growth dynamics of *B. subtilis* in the 384-well plate are noticeably different with a slower growth rate and lower maximum OD compared to the other graphs (**Figure 2A)**. We saw this consistently, highlighting the importance of constructing separate standard curves for each plate type. To calculate the standard curve, lag time is extracted from the growth curve dataset and plotted against the number of cells in that well as determined by CFU counting (**Figure 2B**). Using a simple linear regression, the slope of the standard line can be used in downstream experiments to estimate cell counts.

**Figure 2:**
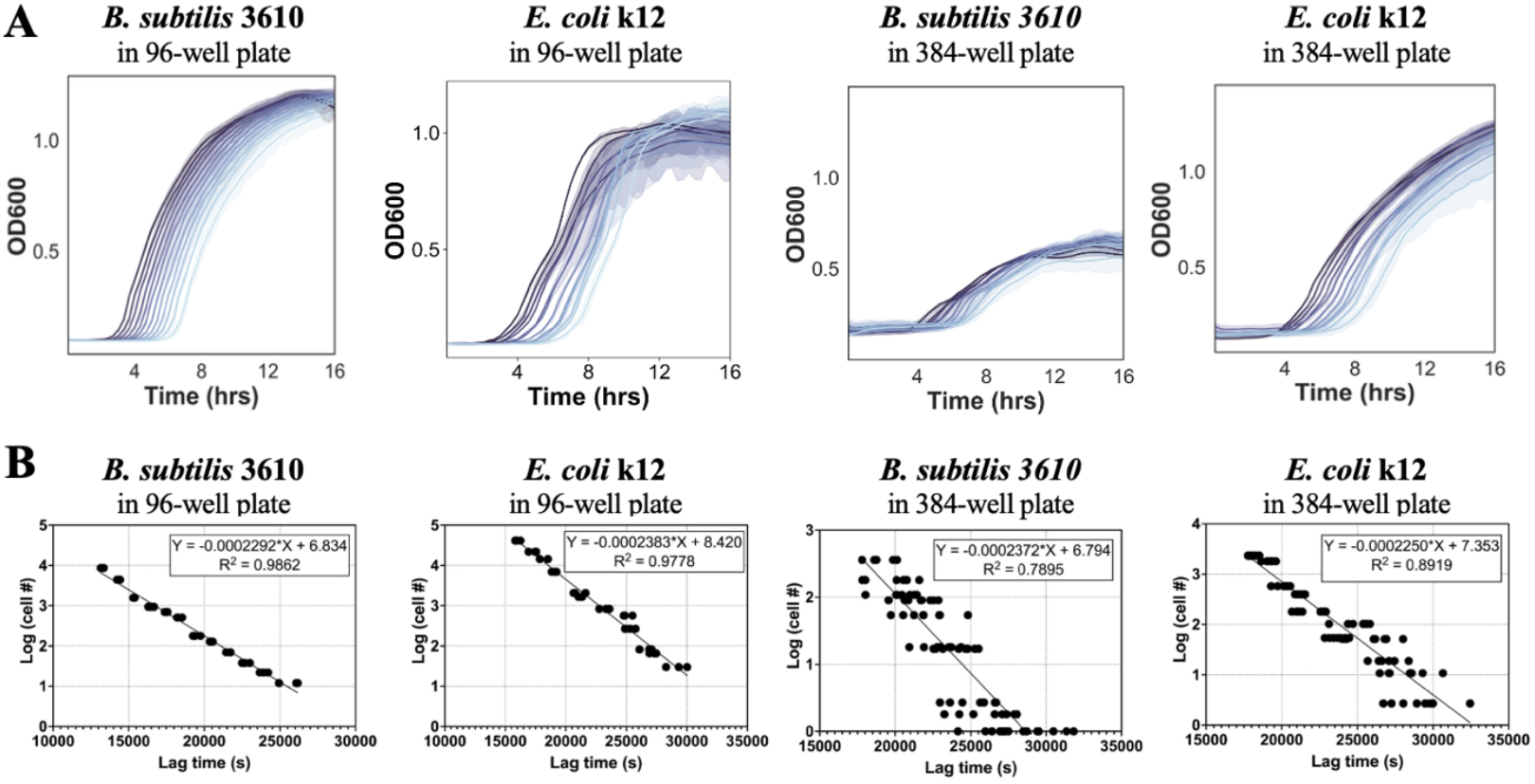
Constructing Standard Curves. Standard curves were constructed for each species in both 96 and 384 well plates by plotting the lag time(s) of each well by the number of cells in each well. **A**. Growth curves for the two-fold serial dilutions of *Bacillus subtilis* 3610 and *Escherichia coli* K12 in 96-well plates and 384-well plates. Growth curves are colored by dilution with the wells with the highest concentration of bacterial cells shown in the darkest shade. **B**. Standard curves are calculated from growth curves by plotting number of cells in each well against lag time. Equations for the best fit line and R^2^ value are shown in the figure.

### 96-well Capillary Assay

The concentration of a chemoeffector at which no distinguishable difference from background can be measured is called the threshold concentration^6^. We were interested in how the sensitivity of our high-throughput method compares to the sensitivity of traditional capillary tube assays for *B. subtilis* and *E. coli*. For *B. subtilis* the threshold concentrations for aspartate and serine were found to be 3×10^−5^ M and 6×10^−7^ M, respectively^7^. The threshold concentration for *E. coli* towards aspartate and serine was identified to be 6×10^−8^ M and 3×10^−7^ M, respectively^6^. To assess the sensitivity of our assay, we made 10-fold dilutions of serine and aspartate stock solutions and tested whether we could observe a difference between the number of cells in our control condition (0 M) compared to the different dilutions.

All concentrations of aspartate tested for *B. subtilis* separated from the 0 M control growth curve (**Figure 3A**). Cell counts remained low for *B. subtilis* towards aspartate. However, all dilutions of aspartate tested had average cell counts greater than the control data with averages of 16.49, 4.88, 3.81, 2.74, 1.64, 1.92, 1.03, 0.86 for aspartate concentrations of 10^−1^ M, 10^−2^ M, 10^−3^ M, 10^−4^ M, 10^−5^ M, 10^−6^ M, 10^−7^ M, 0 M, respectively (**Figure 3B**). The chemotactic index for the aspartate dilutions also highlights this decreasing trend with 10^−1^M aspartate having the highest CI value of 0.95 and 10^−7^ M having a CI value of 0.55, all other dilutions leading to CI values in between (**Figure 3C**). Aspartate concentrations equal to or greater than 10^−6^ M have chemotactic indexes above 0.6, which indicates that they are attractants. Based on these data, we can conclude that the threshold concentration of aspartate chemoattraction in *B. subtilis* in our assay is 10^−6^ M.

**Figure 3:**
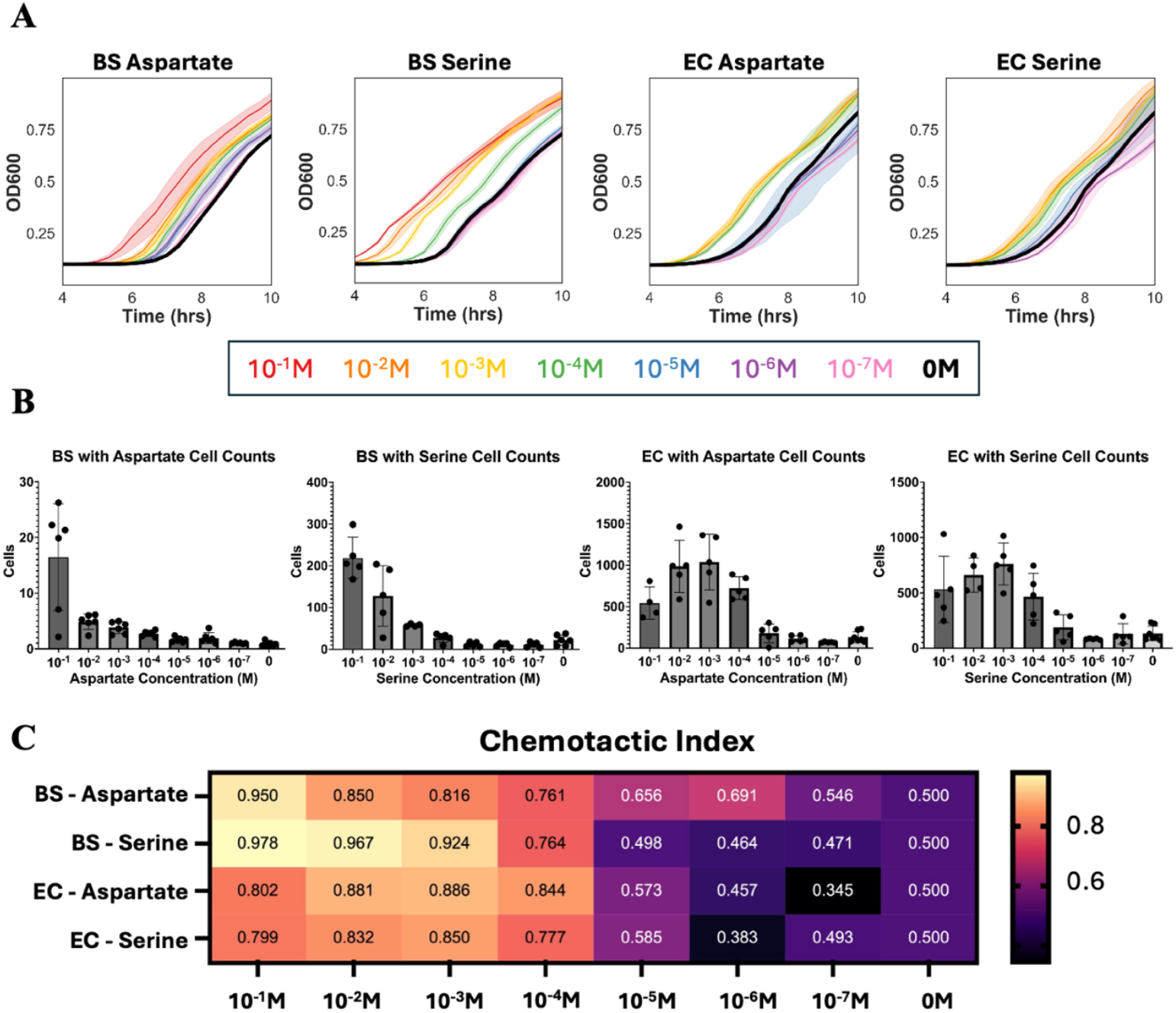
96-well plate threshold concentrations. **A**. Growth curves data from *B. subtilis* (BS) and *E. coli* (EC) chemotaxis experiments in 96-well plates. Growth curves are color coded by the concentration of chemoeffector used in the sample. A legend for the color can be found below the graphs. **B**. Calculated cell counts for *B. subtilis* and *E. coli* capillary tubes containing various concentrations of chemoeffectors. Cell counts are calculated from the growth curve data using the appropriate standard curve. **C**. Heat map showing the Chemotactic Index (CI) calculated for each condition. CI = T / (T+C) where T is the number of cells in the chemoeffector capillary and C is the number of cells in the control capillary. CI > 0.6 is considered an attractant.

In addition to aspartate, we also tested serine and found *B. subtilis* was highly responsive to concentrations of serine above 10^−4^ M. Concentrations below 10^−4^ M did not separate from the control growth curve (**Figure 3A**) and had average cell counts equal to or slightly lower than the control condition (**Figure 3B**). For the higher concentrations, cell counts were much higher than the cell counts for aspartate with averages of 119.02, 76.71, 32.12, 8.50, 2.60, 2.27, 2.34, 2.63 for serine concentrations of 10^−1^ M, 10^−2^ M, 10^−3^ M, 10^−4^ M, 10^−5^ M, 10^−6^ M, 10^−7^ M, 0 M, respectively (**Figure 3B**). From the chemotactic indexes shown in **Figure 3C**, concentrations greater than or equal to 10^−4^ M serine can be considered attractant in our assay with CI values ranging from 0.764 to 0.978. Based on these data, we can conclude that the threshold concentration of serine chemoattraction in *B. subtilis* in our assay is 10^−4^ M.

For *E. coli*, concentrations of aspartate ranging from 10^−1^ M to 10^−4^ M yielded similar results with overlapping curves **(Figure 3A)**. Unlike *B. subtilis*, for *E. coli* the peak response was at 10^−3^ M rather than 10^−1^ M with an average of 541.56, 989.44, 1039.36, 732.15 cells for aspartate concentrations of 10^−1^ M, 10^−2^ M, 10^−3^ M, 10^−4^ M, respectively. After 10^−4^ M, there is a large decline in average cell counts with 179.42, 112.78, 70.62, 133.85 cells in capillary tubes with aspartate concentrations of 10^−5^ M, 10^−6^ M, 10^−7^ M, 0 M, respectively (**Figure 3B**). Therefore, for these measurements 10^−4^ M aspartate was found to be the threshold concentration for *E. coli*. This is also corroborated by the chemotactic index values **Figure 3C** which fall between 0.802 and 0.886 indicating that these concentrations are effective chemoattractants.

Trends for *E. coli* chemotaxis towards serine are similar to the *E. coli* aspartate data. The threshold concentration of *E. coli* for chemotaxis towards serine is also 10^−4^ M with concentrations below this resulting in overlapping growth curves with the motility media control **(Figure 3A)**. Like aspartate, responses to all concentrations of serine tested greater than 10^−4^ M were similar with a peak response at 10^−3^ M **(Figure 3B)**. Concentrations above 10^−4^ M had similar cell counts with averages of 531.90, 661.8, 761.06, 465.84, 188.88, 83.23, 130.38, 133.85, for serine concentrations of 10^−5^ M, 10^−6^ M, 10^−7^ M, 0 M, respectively **(Figure 3B)**. Concentrations of serine above 10^−3^ M had CI values ranging from 0.777 to 0.850 indicating attractants, while concentrations below this threshold had CI values ranging from 0.383 to 0.585 indicating no attraction.

### 384-well Capillary Assay

After optimizing this method using a 96-well plate format, we aimed to further increase the throughput by expanding to 384-well plates. To do this, we 3D printed a new capillary tube holder designed to accommodate the dimensions of Nunc™ 384-Well plates. We then validated the 384-well assay and investigated its sensitivity by repeating experiments done in the 96-well plates and comparing results. As seen in the 384-well standard curves, *B. subtilis* had slightly impacted growth dynamics, with a slower growth rate and lower max OD when compared to *B. subtilis* grown in a 96-well plate (**Figure 4A**). This was not the case for *E. coli* which had a fast growth rate and max OD comparable to what is seen in growth in 96-well plates (**Figure 4A**).

**Figure 4:**
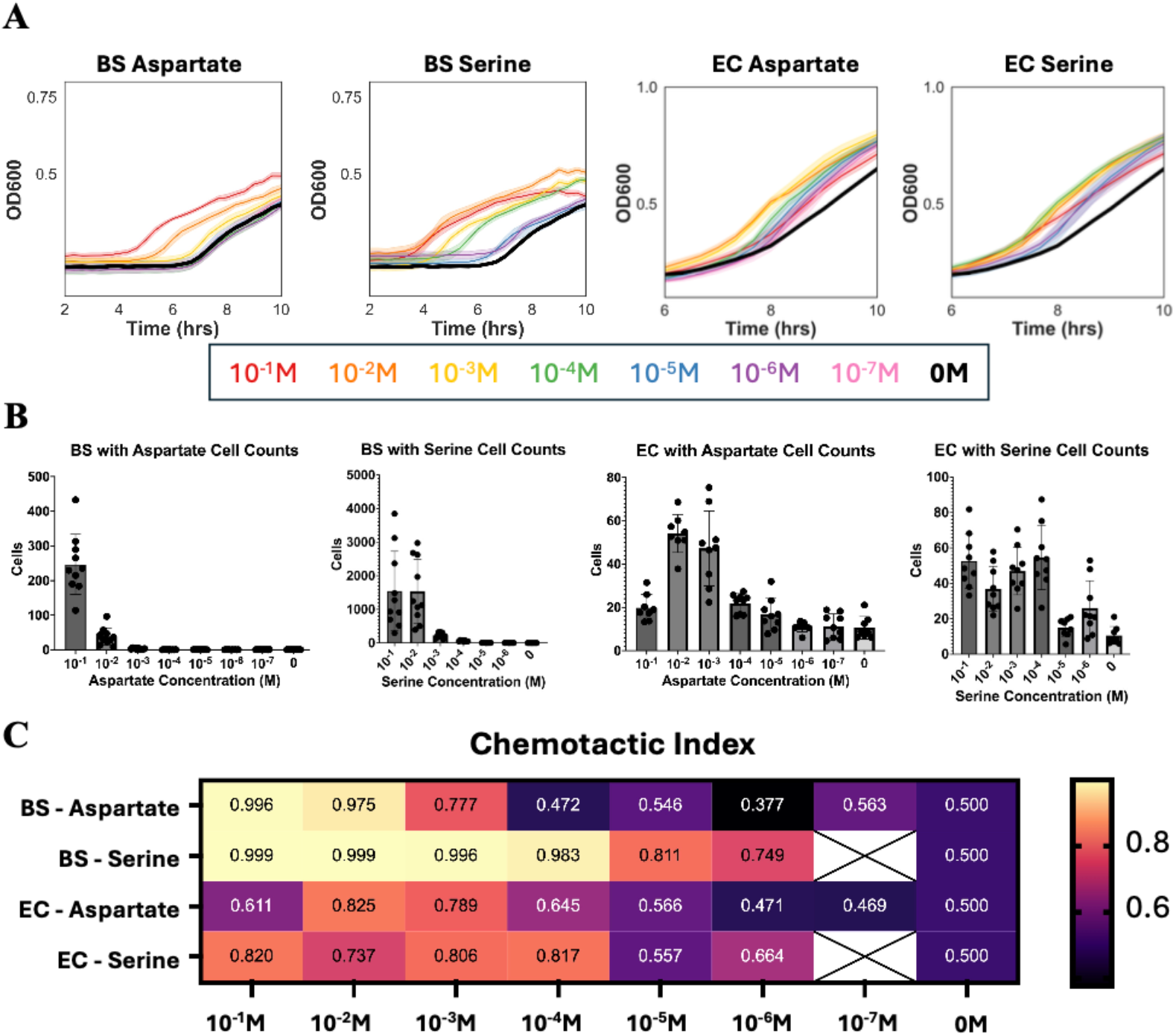
384-well plate threshold concentrations. **A**. Growth curve data from *B. subtilis* (BS) and *E. coli* (EC) chemotaxis experiments in 384-well plates. Growth curves are color coded by the concentration of chemoeffector used in the sample. A legend for the color can be found below the graphs. **B**. Calculated cell counts for *B. subtilis* and *E. coli* capillary tubes containing various concentrations of chemoeffectors. Cell counts are calculated from the growth curve data using standard curves. **C**. Heat map showing the Chemotactic Index (CI) calculated for each condition. CI = T / (T+C) where T is the number of cells in the chemoeffector capillary and C is the number of cells in the control capillary. CI > 0.6 is considered an attractant.

The 384-well plate successfully detected chemoattraction in all cases, where the concentration of the attractant was between 10^−2^ M and 10^−3^ M. *B. subtilis* growth curves for different concentrations of aspartate above 10^−3^ M separated visibly on the growth curve, whereas for serine, all concentrations above 10^−6^ M separated visibly on the growth curve suggesting a lower threshold concentration for serine compared to aspartate (**Figure 4A**). In general, serine elicited a stronger response from *B. subtilis* than aspartate with the highest cell counts for serine capillaries in the 10^−1,^ M dilution with 1545.32 cells, while the highest response for aspartate was in the 10^−1^ M dilution with 246.83 cells (**Figure 4B**). The stronger response of *B. subtilis* towards for serine was also reflected in the CI values for the lower concentrations, which were all greater than 0.7 indicting a chemoattraction (**Figure 4C**). For aspartate, the threshold was higher at 10^−3^ M with a CI value of 0.77 (**Figure 4C**).

*E. coli* data collected using the 384-well plate closely resembled the data collected using our 96-well plate with threshold concentrations for both serine and aspartate at 10^−4^ M. For aspartate, *E. coli* peak response was found to be 10^−2^ M, while for serine it was 10^−1^ M. *E. coli* growth curves for the dilutions of aspartate and serine separated from the control but were more overlapping than the *B. subtilis* curves. This indicates a non-linear response of *E. coli* to the different concentrations of chemoeffectors (**Figure 4A**). *E. coli* showed similar magnitudes of response to both aspartate and serine with peak cells counts at 54.6 and 52.9 for aspartate and serine, respectively (**Figure 4B**). CI values for *E. coli* indicate that all concentrations greater than or equal to 10^−4^ M for both chemoeffectors are chemoattractants (**Figure 4C**).

## Discussion

In this study, we optimized a high-throughput capillary tube-based chemotaxis assay to allow for rapid screening of chemoattractants. We demonstrated that the system is effective for studying both gram-negative and gram-positive organisms and that the assay is functional in both a 96- and 384-well format. This greatly expands on the current capabilities of the field, describing for the first time a method to simultaneously run up to 384 capillary tube assays, enabling researchers to expand the libraries being tested when describing chemotactic profiles of bacteria of interest.

The throughput of the assay relies upon the relationship between lag time and starting cell concentration. This allows for the use of growth curves in plate readers for data collection and dramatically reduces the workload and user-error associated with traditional quantification strategies such as CFU counting. The assay can be adapted to work with any plate layout that can be read using a plate reader. We therefore think that the throughput could be further expanded to include 1536-well plates. The reliance on plate readers for quantification, however, does introduce some limitations. It requires that the bacteria being studied is compatible with plate-based growth in a manner in which the relationship between lag time and OD is maintained. Additionally, sources of error inherent to the plate reader become relevant considerations here. This includes the influence of evaporation, which may cause edge effects that may become more prominent when using smaller well volumes or running growth curves for prolonged periods of time. This may be avoided by filling edge wells with water rather than samples.

The sensitivity of our assay was investigated for both the 96- and 384-well layouts. We found that for the 96-well plate, threshold concentration for *B. subtilis* towards aspartate and serine was 10^−6^ M and 10^−4^ M, respectively. This is aligned with previously reported threshold concentrations. For the 384-well plates, the threshold concentrations were 10^−3^ M and 10^−7^ M, for aspartate and serine, respectively. Discrepancies in these values may be due to the differences in the well plates, as *B. subtilis* did exhibit altered growth behavior in the 384-well plate, which reduced the fit of our standard curve resulting in a lower R-squared value compared to the 96-well plate. This likely impacted the accuracy of our cell count calculations for the 384-well plate, suggesting that in some cases, there may be a trade-off between throughput and sensitivity if lower chemoattractant concentrations are being studied.

For *E. coli*, the threshold concentrations for aspartate and serine were consistently at 10^−4^ M irrespective of whether the 96- or 384-well plate was used. These threshold values for *E. coli* are higher than what was previously reported for manual capillary tube. However, we show that the assay can detect chemoattraction towards concentrations that span many magnitudes of order, which should be sufficient for most high-throughput screening purposes. Additional limitations to this assay exist that are inherent to all capillary tube assays. This includes the requirement of swimming motility and the inability to draw conclusions about chemorepulsion. Additional work is needed to develop a high-throughput assay to screen bacterial chemorepellents and bacteria which use twitching- or swarming-based motility.

In summary, here, we have described a novel 3D-printed capillary tube holder that can be used to conduct up to 384 individual capillary tube assays simultaneously. We show our system is effective in both the 96-well and 384-well formats and that it is sufficiently sensitive to detect changes in chemotactic behavior against a gradient of chemoeffector concentrations in both gram-negative and gram-positive bacteria. This method greatly reduces the time required to screen large libraries of chemoeffectors and will enable the generation of larger datasets characterizing bacteria chemotactic behavior.

## Materials & Methods

### Materials

Fisherbrand™ 96-Well plates (Cat. No. FB012931) and Nunc™ 384-Well plates, (Cat. No. 242757) were used for our assays along with 1µL capillary tubes from Drummond (Cat. No. #1-000-0010) and TempPlate XP sealing film (USA SCIENTIFIC #2972-2100).

### Designing and printing 3D capillary tube holder

Designs were created using OnShape and printed at the Caltech TechHub on a Bambu X1 Carbon printer using TPU filament (ZIRO TPU Filament 1.75 mm, Shore 95A). The .stl file was sliced for printing using the Orca slicer software, 0.20 mm standard settings were used with slight modifications: Bed Type = “Smooth High Temp Plate” and Filament type = “Generic TPU”. For new prints, we recommend going through each hole with a 20-gauge needle (or needle with an outer diameter closest to the desired hole diameter) before trying to insert the capillaries. The capillaries should fit tightly but be able to be inserted and removed without breaking. For optimization a calibration print is including in **Supplemental Files S1** which can be used to determine the optimal capillary hole diameter based on the print conditions. Print files for the calibration piece, 96-well capillary holder, and 384-well capillary holder can be found in **Supplemental Files S1-S3**.

### Testing capillary expulsion speed

Capillary tube holders were filled with empty capillary tubes and placed atop a well plate containing food coloring. The capillaries were allowed to fill with food coloring before moving to an empty plate. The capillary tube holder on the empty plate was placed in an Eppendorf Centrifuge 5910 Ri with a second 3D-printed capillary tube holder acting as a balance. Various spin settings were tested for complete expulsion which was confirmed by observing remaining food coloring in the capillary tubes after spinning.

### Preparing bacteria cells for experiments

*E. coli* k12 (ATCC25404) and *B. subtilis* 3610 (ATCC6051) were inoculated into LB media and allowed to incubate at 37°C shaking at 250 rpm for 16-18 hours. Overnight cultures were diluted approximately 100-fold in fresh LB media and grown at 37°C shaking at 250 rpm until mid-exponential phase (OD_600_ of 0.35-0.6) to ensure high motility and cell viability before assay preparation. The volume of each culture needed to obtain 1.5 mL of cells at an OD_600_ of 1.0 was then calculated from the measured OD_600_. The corresponding volume of overnight culture was centrifuged for 5 minutes at 1700 × *g*, and the supernatant was removed. The pellets were washed with Motility Media (10 mM Potassium Phosphate buffer, pH 7; 0.1 mM EDTA; 0.05% glycerol) and resuspended by gentle inversion to avoid disrupting motility machinery. This wash step was repeated again under the same conditions. Following the second wash, the cultures were centrifuged with the same parameters and resuspended in 1 mL of motility media. OD_600_ was checked following washing, and the cell suspensions were diluted to the desired OD_600_.

### Constructing standard curves

*E. coli* and *B. subtilis* cultures were prepared and washed as described in *Preparing cells for experiments*. After the final resuspension, the cultures were diluted with LB to an OD_600_ of 10^−4^ from which 11 two-fold serial dilutions in LB were made. From the serially diluted cultures, 180 µL was added to each of the designated wells of a 96 or 384 well plate, with at least three replicates for each concentration of cells. The plate was placed in a Tecan Infinite M Nano plate reader for 24 hours at 37°C and with continuous orbital shaking at ~200 rpm, recording OD_600_ at 20-minute intervals. CFU counting of each serially diluted culture was used to determine the number of cells in each well. The standard curves were calculated from the resulting growth curves by plotting the number of cells in each well against lag time (time required for each well to reach OD of 0.3).

### 96-well plate chemotaxis assay

*E. coli* and *B. subtilis* cultures were prepared and washed as explained in *Preparing cells for experiments*. The cell suspensions were diluted with motility media to a final OD_600_ of 0.01. A total of 180μL of each diluted culture was then added to the designated wells of a 96-well plate. The plate was then allowed to incubate statically at 37°C for 30 minutes to allow cells to recover after washing.

To prepare the chemoeffector plate, 180 μL of seven ten-fold serially diluted concentrations (10^−1^ M, 10^−2^ M, 10^−3^ M, 10^−4^ M, 10^−5^ M, 10^−6^ M, and 10^−7^ M) of serine and aspartate were added to their designated wells, with motility media in control wells. A wash plate was prepared by adding 200 μL of sterile ddH_2_O water to each well of a third 96-well plate. After the bacteria plate was finished recovering, the capillary holder containing 1 µL capillaries in each opening was placed on top of the chemoeffector plate. The capillary holder was left on for 2 minutes to ensure each of the capillaries filled. The top of the capillary tube holder was sealed tightly with plate film, transferred to the wash plate, and dipped up and down three times to remove excess chemoeffector solution on the outside of the capillaries. The holder was then transferred to the bacterial plate and incubated at room temperature for 45 min. Following incubation, the capillary tube holder containing the capillaries was placed on a blank plate and centrifuged for two minutes in an Eppendorf Centrifuge 5910 Ri at 1000×*g* to expel their contents. After centrifugation the capillary tube holder was removed and total of 180 μL of LB media was added to each well, and the plate was placed in a Tecan Infinite M Nano plate reader for 24 hours at 37°C and with continuous orbital shaking at ~200 rpm, recording OD_600_ at 20-minute intervals.

### 384-well plate chemotaxis assay

*E. coli* and *B. subtilis* cultures were prepared and washed as explained in *Preparing cells for experiments*. The cell suspensions were diluted with motility media to a final OD_600_ of 0.01 With each of the diluted cultures, 60 µL was added to the designated wells of a 384-well plate. The plate was then allowed to incubate statically at 37°C for 30 minutes to allow cells to recover after washing.

The chemoeffector plate was then prepared by adding 60 μL of seven ten-fold serially diluted concentrations (10^−1^ M, 10^−2^ M, 10^−3^ M, 10^−4^ M, 10^−5^ M, 10^−6^ M, and 10^−7^ M) of serine and aspartate to their designated wells, with motility media in control wells. A wash plate was prepared by adding 80 μL of sterile ddH_2_O water to each well of a third 384-well plate. The steps for filling, washing, incubating, and expelling the capillaries were performed as described for the *96-well plate chemotaxis assay*, except that a capillary holder with 384 openings was used in place of the 96-well holder. After the capillary contents were expelled, a total of 90 μL of LB medium was added to each well, and the plate was placed in a Tecan Infinite M Nano plate reader for 24 hours at 37°C and with continuous orbital shaking at ~200 rpm, recording OD_600_ at 20-minute intervals.

## Author contributions

Conceptualization: CB & GSD

Methodology: CB, EG, SE, & GSD

Data Acquisition: CB, EG, & SE

Supervision: CB & GSD

Writing & drafting: CB, EG & GSD

Funding Acquisition: CB & GSD

## Acknowledgements

We would like to thank Ian Roberts and Arleen Hom at the Caltech TechHub for their assistance with 3D printing.

